# Screening coffee genotypes for *Cercospora coffeicola* resistance in Brazil

**DOI:** 10.1101/2021.10.07.463543

**Authors:** Juliana Barros Ramos, Mario Lucio Vilela de Resende, Deila Magna dos Santos Botelho, Renata Cristina Martins Pereira, Tharyn Reichel, André Augusto Ferreira Balieiro, Gustavo Pucci Botega, Juliana Costa de Rezende Abrahão

## Abstract

Several efforts have been made by many researchers worldwide to develop coffee plants resistant to different *Cercospora* species; however, studies concerning *C. coffeicola*, specifically, are still incipient. In the present study, a blend of strains from this pathogen was inoculated into 18 Brazilian commercial cultivars, a coffee clone of Arabica, as well as into 41 accessions from the Germplasm Collection of Minas Gerais, to evaluate the genetic resistance ability within the population and select superior genotypes for the breeding program. After predicting genotypic values of the evaluated material, the most efficient way to select genotypes based on the data of severity to brown eye spot (BES) was also examined. Moreover, the action of defense mechanisms against *C. coffeicola* attacks was investigated by assessing the levels of total soluble phenolic compounds and soluble lignin in contrasting genotypes regarding disease susceptibility. Based on the results, the accession MG 1207 Sumatra demonstrated an intrinsic genetic capacity to maintain low levels of severity to brown eye spot. This genotype can thus substantially contribute to the development of new cultivars, which may lead to reduced use of pesticides. This study also evidenced that four evaluations of severity is enough to reach accuracy and efficiency for the severity of BES, thus providing expressive genetic gains. Finally, it is suggested that the levels of lignin and phenolic compounds are not associated with the resistance of coffee genotypes to brown eye spot.

## Introduction

Cercosporiosis or ‘brown eye spot’ (BES), a disease in which the etiologic agent is the necrotrophic fungus *Cercospora coffeicola* Berk. & Cooke, has been pointed out as of great economic importance for coffee growers. This disease causes defoliation, stimulates maturation, and increases the number of empty beans, besides intensifying the pulp adherence to the endocarp - which makes it difficult to de-pulping - thereby affecting the quality of the produced coffee [1]. In the absence of control management, losses can reach 30% [2]. In Brazil, increases in the incidence of brown eye spot in coffee fields by the late 2000’s coexisted with the expansion of coffee fields to other regions with different environmental conditions. Moreover, the cultivation of new coffee varieties and the use of different cultural practices have benefited the development of this pathogen under a climate change condition [3].

Disease management has been commonly accomplished by the application of chemical products coupled with an accurate water supply and plant nutrition [4]. To date, as the physiology of plant responses underlying genetic resistance is quite complex, few studies have been carried out on brown eye spot resistance in coffee species [5, 6, 7]. In this context, Dell’Acqua et al. [5] and Patricio et al. [6] used the phenotypic data from brown eye spot severity evaluation - in a greenhouse condition - to carry out the selection based on a test of means. These authors considered the studied genotypes as fixed effects, without, however, estimating the genetic variance and heritability of the phenotypic variation. This strategy may not be effective in crop breeding programs that aim at developing materials resistant to BES as it does not assess the potential effects of selection on the studied characters.

To defend themselves against pathogen attacks, plants have a set of defense mechanisms, which can be either constitutively structural or biochemical; as well as a defense system responsive to infection occurrences [8]. Among the induced mechanisms, the production of phenolic compounds and lignin is commonly observed during defense reactions, which are produced via metabolism of phenylpropanoids [9], thereby providing higher resistance to the plant cell wall during pathogen attack. Hence, understanding how *C. coffeicola* affects biochemical aspects of the plant cell can base a further mechanism to control brown eye spot.

The first objective of this study was to select superior *Coffea arabica* genotypes concerning resistance to BES. The possibility of selecting coffee plants with higher resistance to this disease is extremely important, but the success of this strategy depends on the existence of genetic variation within studied characteristics, as well as a high heritability rate. The second objective was to better understand the action of defense mechanisms against the attack of the pathogen *C. coffeicola*, by quantifying total soluble phenolic compounds and soluble lignin in contrasting genotypes regarding disease susceptibility. It is worth mentioning that no studies were found relating the concentration of such compounds in coffee genotypes with resistance to brown eye spot.

## Material and methods

### Brown eye spot severity

The experiment was carried out in greenhouse conditions at the Department of Phytopathology of the Universidade Federal de Lavras - UFLA, Brazil. Coffee seedlings of 18 commercial cultivars, an Arabica coffee clone (Siriema 312), and 41 accessions from the Germplasm collection of the Agricultural Research Corporation of the State of Minas Gerais (EPAMIG) in Patrocínio, MG, were evaluated for resistance to brown eye spot (Table S1). Accessions were selected according to their characteristics of yield, drink quality and/or resistance to other diseases of economic importance.

Seeds were sown in 5L plastic trays containing autoclaved sand. The germination chamber was adjusted to 30°C and 80% relative humidity. The seedlings were transplanted into punched-black polyethylene pots (0.11 x 0.20 m) at the phase of cotyledons. The substrate consisted of 300L of cattle manure and 700L of soil mix extracted from the 0.4 to 0.8m layer of Dystrophic Red Latosol and fertilized with 5kg of simple superphosphate and 500g of potassium chloride. Sprinkler irrigation was performed on all genotypes, during mornings and afternoons, in a manner that the substrate was kept at field capacity.

The seedlings were inoculated at the stage of three pairs of true leaves. The inoculum consisted of a blend of different isolates of *C. coffeicola* obtained from coffee leaves with symptoms of brown eye spot collected in the municipalities of Marechal Floriano (ES), Cachoeirinha, Ervália, Lavras and Patrocínio (MG). The objective of using different isolates was to include a higher variability of the fungus [1, 7].

The sporulation of the isolates was performed as described by Souza et al. [10], with adaptations. Eight mycelial discs (6mm in diameter) were removed from the colony borders (at the 15^th^ days of growth) over different isolates of *C. coffeicola.* Discs were macerated in 400μL of sterile distilled water. The macerated mycelium from each isolate was placed in Erlenmeyer flasks containing 20mL of liquid V8 culture medium (100mL of V8 in 900mL of distilled water) under shake at 100 rpm for 12 d at room temperature. The liquid containing the mycelium was transferred to plates with a water-agar medium. The plates were kept in a BOD incubator (Bio-Oxygen Demand), with a photoperiod of 12 h at 25°C. After culture medium dehydration (approximately 5 days of incubation), 10 ml of sterile water was added to each plate and the conidia were removed with a Drigalski spatula. The suspension was filtered with sterile gauze to remove residues and the conidia were subsequently quantified in a Neubauer chamber. The suspension used for inoculation was adjusted to 5 x 10^4^ conidia mL^-1^ and sprayed on the abaxial side of the leaves of all seedlings, by using a manual sprayer. Afterward, the seedlings were placed in a humid chamber for 72 h.

Temperature and relative humidity data were collected over the experiment with a Datalogger HT-500, Instrutherm^®^. Weekly assessments concerning disease severity were performed on the first two pairs of true leaves during five weeks, starting from the onset of symptoms (about 15 d after inoculation). The severity of brown eye spot in different coffee genotypes was quantified using a diagrammatic scale with six classes of the proportion in the infected area by brown eye spot [11]: class 1: 0.1-3.0%; class 2: 3.1-6.0%; class 3: 6.1-12.0%; class 4: 12.1-18.0%, class 5: 18.01-30.0% and class 6: 30.1-50.0%. The experiment was repeated twice. The distribution of the phenotypic segregation was also evaluated in each genotype within the six classes described. Based on this assessment, the resistance level was determined: class 1, as resistant (R); class 2, as partially resistant (PR); class 3, as moderately susceptible (MS); class 4, as susceptible (S); classes 5 and 6, as highly susceptible (AS).

### Total soluble phenolic compounds and soluble lignin

Both total soluble phenolic compounds and soluble lignin levels were quantified in leaves of MG 1207 genotype (presenting low disease severity in the present study) and MG 0291 and Catuaí Vermelho IAC 144 genotypes (presenting high disease severity). The second one was used as a control of susceptibility, according to Patricio et al. [6] and Botelho et al. [7].

The samples consisted of 2nd and 3rd pairs of fully expanded leaves, collected at 24, 120, 240, 480, and 720 h after inoculation (hai) of *C. coffeicola*. Samples of non-inoculated plants with the pathogen were also collected at t 24 and 720h to confirm whether the inoculation influences the levels of total soluble phenolic compounds and soluble lignin. After collection, the samples were immediately stored in liquid nitrogen and then in an ultra-freezer until sample processing.

The macerated samples were lyophilized and approximately 30 mg of the material were homogenized in 80% methanol. The solution was centrifuged at room temperature at 14000 rpm for 5 min. The supernatant and the precipitate were used to quantify total soluble phenolic compounds and soluble lignin, respectively.

The levels of total soluble phenolic compounds were determined as described by Spanos and Wrolstad [12], with modifications. The supernatant was homogenized with Folin-Ciocalteau reagent at 0.25N, 1M Na_2_CO_3_, and distilled water. The reaction was standardized in 200 μL and quantified in a spectrophotometer at 725nm. Based on the standard curve of chlorogenic acid, the levels of total soluble phenolic compounds were calculated.

Lignin was quantified as proposed for Doster and Bostock [13]. The precipitate, which was homogenized in 80% methanol and centrifuged as described for phenolic compounds. The contents were evaporated in an oven at 45°C overnight and mixed with thioglycolic acid and 2 M HCl (ratio 1:10) in a water bath at 100°C for 4h. After centrifugation and solubilization in 0.5 M NaOH, the supernatant was homogenized with HCl P.A., and kept at 4°C for 4 h before centrifugation. The precipitate was homogenized in 0.5M NaOH. The aliquot of 200 μL of this solution was used for the reaction, and the absorbance was assessed in a PowerWave XS microplate spectrophotometer (Biotek^®^) at 280 nm. Based on a standard curve of lignin, the soluble lignin content was subsequently estimated. The quantification of total soluble phenolic compounds and soluble lignin contents was performed in triplicates.

### Statistical analysis

The evaluate of the disease severity of brown eye spot in the coffee genotypes, the experiments were conducted in a complete randomized block design, with 60 treatments (genotypes), eight replications, and five times of evaluation. Two coffee seedling was considered as experimental unit. The mixed model analysis was accomplished by the Selegen REML/BLUP software [14]. Regarding a completely randomized design, the following equation was used: *y* = *Xm* + *Zg* + *Tp* + *e*, in which *y* is the data vector; *m* is scalar referring to the general average of fixed effect; *g* is the vector of random genetic effects; *p* is the vector of the random-effects of blocks within replications and experiment; and *e* the vector of random errors. X, Z and T are incidence matrices of referred effects. The variance components were submitted to the likelihood ratio test at 5% probability.

The severity dates were used to calculated the area under the curve of disease progress (AUCDP), as previously proposed by Shaner and Finney [15].

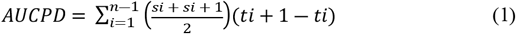

Where: AUCDP = area under the curve of disease progress; Si = disease severity in the time of evaluation; ti = time of evaluation i. The analysis of variance was carried out in a randomized block design with 60 treatments and 8 blocks, with an experimental plot consisting of two plants.

Concerning the analysis of total soluble phenolic compounds and soluble lignin, a completely randomized design was adopted, with three replications and two plants per plot. The analysis of variance was in a 3 x 5 factorial scheme, with three evaluated genotypes and five times of collection, namely 24, 120, 240, 480, and 720 h after inoculation (hai) of *C. coffeicola*. The data were submitted to analysis of variance and the means were compared by Tukey test at 5% probability.

## Results

### Genotype selection

The analysis based on mixed models provided a simultaneous estimation of both individual heritability and plot repeatability (Fig 1, Table S1). The heritability and genetic variation coefficients were significant, thereby indicating the possibility of selection. The selection among families was high (98%) and this should be recommended. The coefficient of permanent effect determination showed a low magnitude (‹0.00), revealing that variations in the environment among measurements do not affect such responses. The overall average of the experiment was over 14.50% of severity (Table S1).

**Fig 1.**
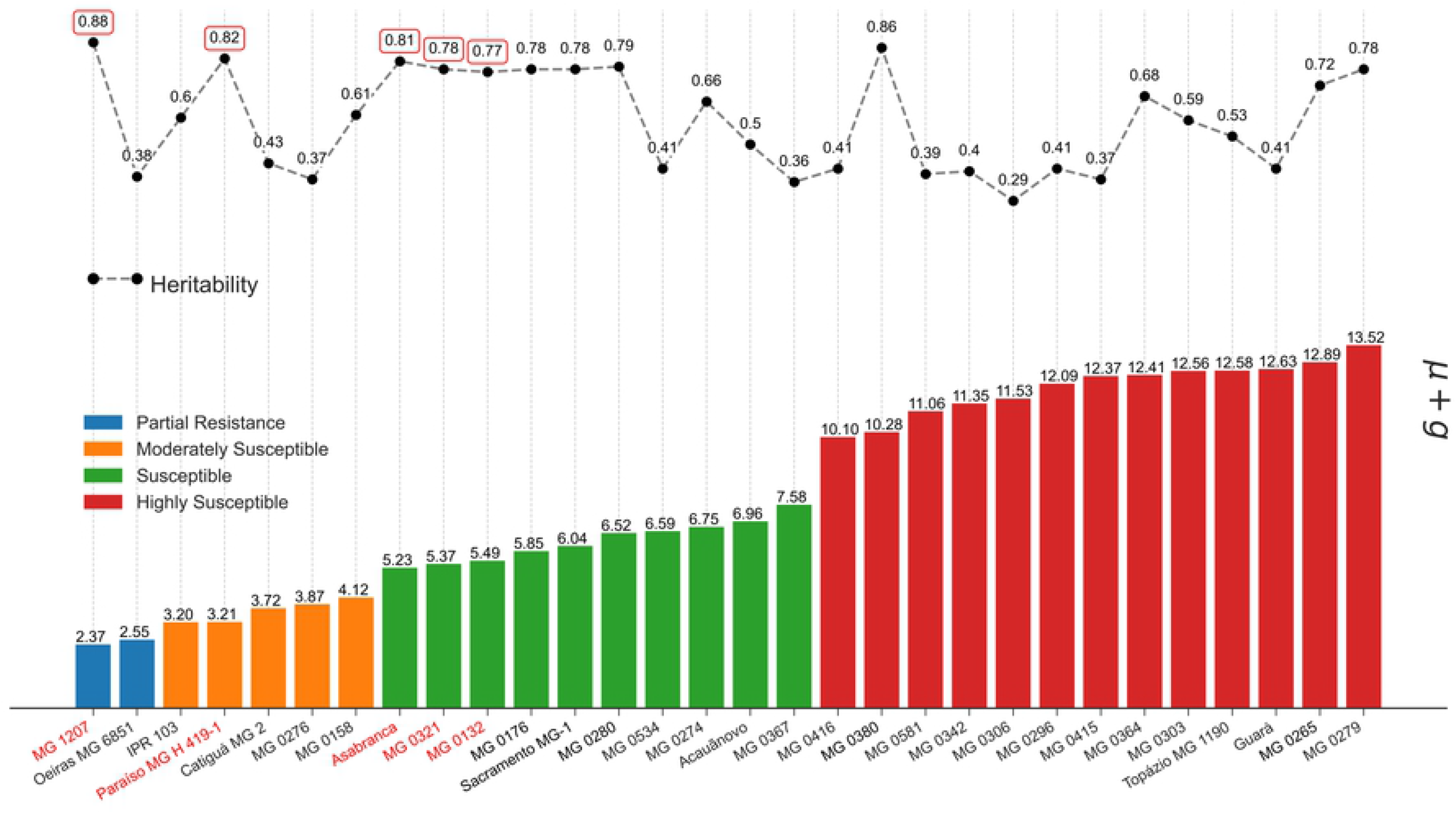
Accession classification based on phenotypic mean, genotypic values (u + g), genotype heritability, calculated on BES severity in coffee seedlings.

The studied population presented a high genotypic variability for the severity of brown eye spot. The predicted additive values ranged between 2.37% (accession MG 1207) and 32.02% (Catuaí Vermelho IAC 144); while the individual heritability showed values between 29 and 88%, among 60 genotypes (Table S1).

Based on the analysis of frequency-severity, it was observed that the genotypes were clustered mainly into 4, 5 and 6 classes (Table 1), thus being considered as susceptible or highly susceptible. The genotypes MG 1207 Sumatra and Oeiras MG 6851 displayed injured leaf area by about 3.1 to 6.0% and were classified as partially resistant (class 2). The accessions MG 0276, MG 0158, and the cultivars Catiguá MG2, Paraíso MG H 419-1 and IPR 103 were considered moderately susceptible, hence class 3. None of the genotypes studied were clustered in the first class (0 to 3% of injured area).

**Table 1.** Average heritability, accuracy and efficiency based on the different number of measurements for the severity of brown eye spot.

| Measurements | 1 | 2 | 3 | 4 | 5 | 6 | 7 | 8 | 9 | 10 |
| --- | --- | --- | --- | --- | --- | --- | --- | --- | --- | --- |
| Heritability | 0.90 | 0.95 | 0.97 | 0.97 | 0.98 | 0.98 | 0.98 | 0.99 | 0.99 | 0.99 |
| Accuracy | 0.95 | 0.97 | 0.98 | 0.99 | 0.99 | 0.99 | 0.99 | 0.99 | 0.99 | 0.99 |
| Efficiency | 1.00 | 1.14 | 1.20 | 1.23 | 1.25 | 1.27 | 1.28 | 1.29 | 1.30 | 1.30 |

Among the studied population, about 38% presented heritability (H) of high magnitude (higher than 70%), namely: MG 1207, MG 0321, MG 0132, MG 0176, MG 0280, MG 0380, MG 0265, MG 0279, MG 0324, MG 0267, MG 0723, MG 0333, MG 0134, MG 0179, MG 0282, MG 0663 and MG 0291 accessions, besides the cultivars Paraíso MG H 419-1, Asabranca, Araponga MG1, Sacramento MG1 and Catucaí Amarelo 2SL and o clone Siriema 312. Within this population, there was a high variation in the severity of brown eye spot, with a phenotypic value ranging from 4.6 to 50.

We selected five genotypes with the highest predicted additive values (between 2.37 and 5.49) and the highest magnitudes of individual heritability (H) (between 77 and 88%), such as the MG 1207, MG 0321, and MG 0132 accessions, and the commercial cultivars Paraiso MG H 419-1 and Asabranca, which presented low magnitude of their genotypic value to reduce the severity, suggesting further probabilities of genetic progress through these evaluations. The selection of five such genotypes resulted in a remarkable selection gain of 70.16% to reduce the severity of brown eye spot.

Based on the phenotypic data, it was observed that the outputs from AUCDP severity for BES presented significant differences among the genotypes. Otherwise, by considering the heritability parameter, the selection via AUCDP highlighted MG 1207, Oeiras MG 6851, IPR 103, Paraíso MG H 419-1 and Catiguá MG 2 genotypes, which showed the lowest disease progress during the evaluated timeframe (Fig 2). Genotypes MG 0291, Catuaí Vermelho IAC 144 and clone Siriema 312 were the most susceptible among them, with a AUCDP of 958.04, 956.24 and 888.08, respectively. By comparing the genotypes that demonstrated the highest and lowest AUCDP, a difference of 901.71% in disease severity was observed.

**Fig 2.**
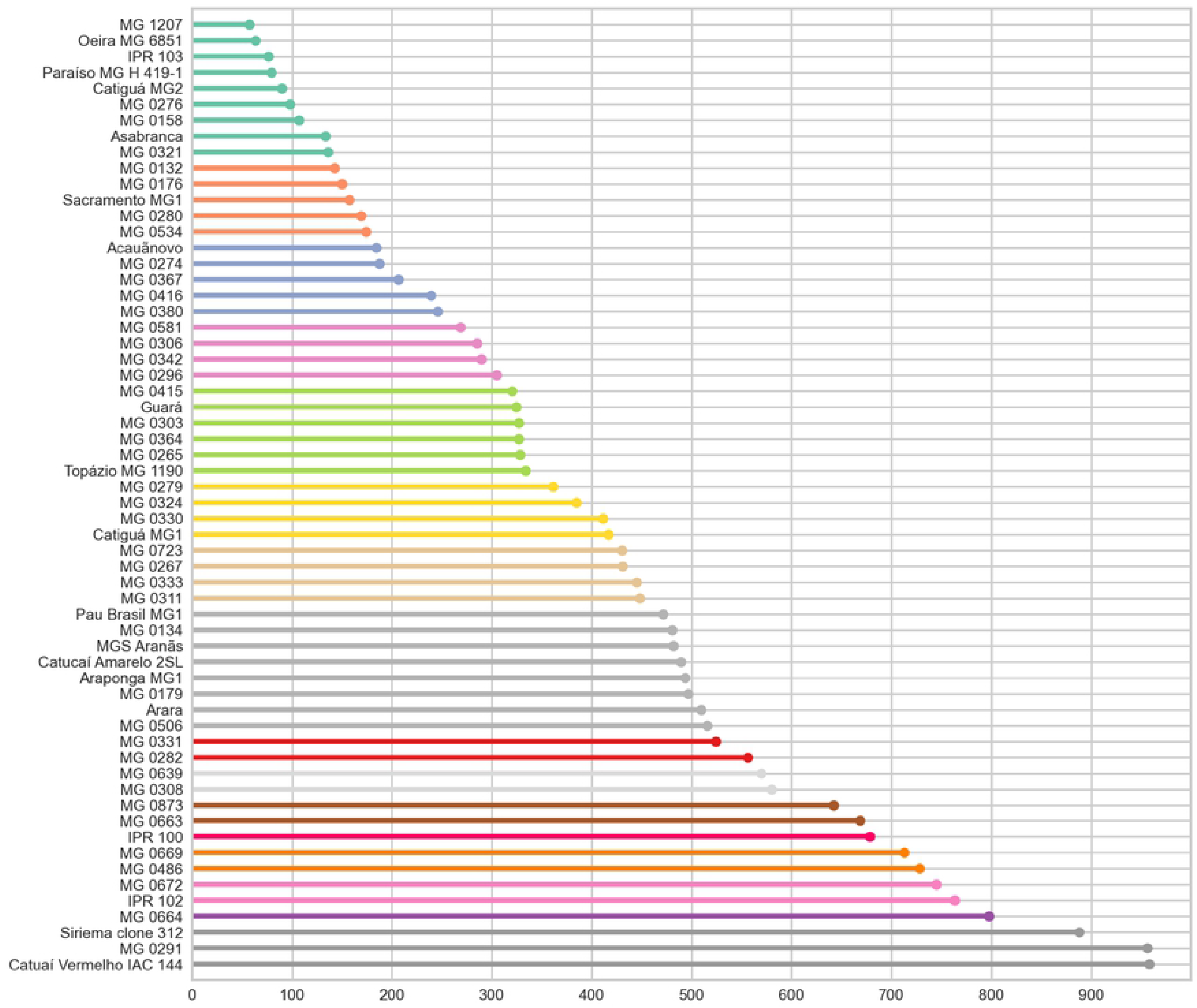
Area under the disease progress curve (AUDPC). *Bars with the same color do not differ from each other by the Scott-Knott test (p≤0.05).

The accuracy can be improved by more rigorous analysis methods, for instance, by increasing the number of measurements per plant. In this context, Table 1 presents the accuracies that would be achieved by using a higher number of measurements. Considering the estimated individual repeatability, it was observed that four measurements lead to accuracies above or close to 99% in the inference about the individuals’ permanent phenotypic values concerning brown eye spot severity.

### Defense mechanisms against *C. coffeicola*

Based on the results of brown eye spot severity, accessions from Sumatra group were selected for quantification of total soluble phenolic compounds and soluble lignin. They were MG 1207, as low severity, and MG 0291, which presented a high severity, being thus placed alongside the control (Catuaí Vermelho IAC 144).

There were no significant differences in total soluble phenolic compounds and soluble lignin levels among studied genotypes, indicating that the action of defense mechanisms against the *C. coffeicola* attack does not differ in terms of disease susceptibility. On the other hand, there was a significant interaction between time of collection for both the contents of phenolic compounds (Fig 3) and soluble lignin (Fig 4).

**Fig 3.**
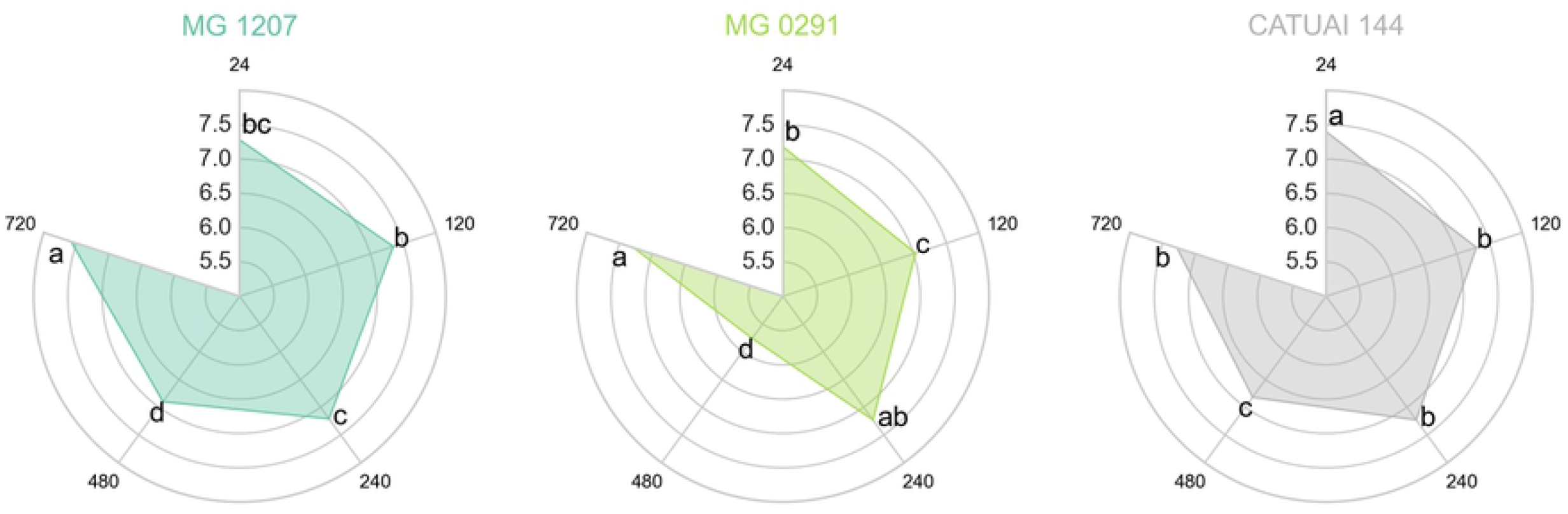
Contents of total soluble phenolic compounds (μL/mg) quantified in leaves of the genotypes MG 1207, MG 0291 and Catuaí Vermelho IAC 144 collected at 24, 120, 240, 480 and 720 h after inoculation of *C. coffeicola*. *Averages followed by the same lowercase letter in the same genotype do not differ by the Tukey test (p≤0.05).

**Fig 4:**
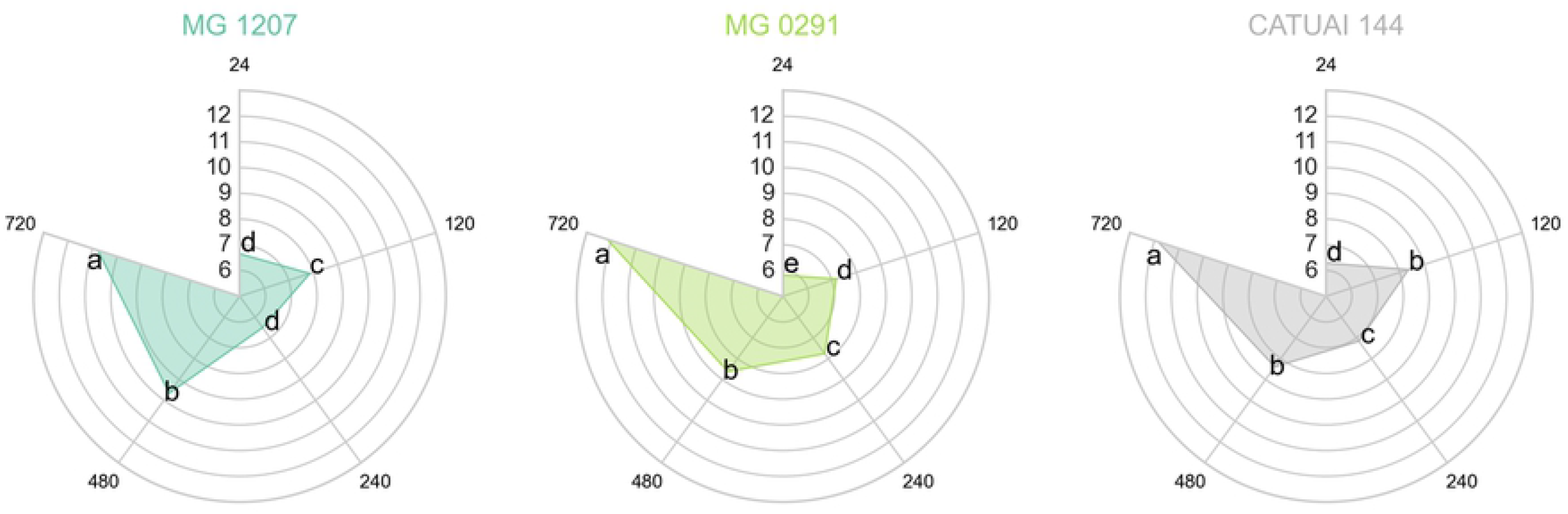
Soluble lignin content (μg/mg) in leaves of MG 1207, MG 0291 and Catuaí Vermelho IAC 144 collected at 24, 120, 240, 480 and 720 h after inoculation of *C. coffeicola*. *Averages followed by the same lowercase letter in the same genotype do not differ by the Tukey test (p≤0.05).

Phenolic compounds content showed variation among the three genotypes over time (Fig 3), with the lowest levels at 480 hai, presenting mean values between 5.76 and 6.90 μL/mg. On the other hand, the highest levels of phenolic compounds were observed at 24 hai in Catuaí Vermelho IAC 144 (7.40 μL/mg) and 720 hai in MG 1207 (7.58 μL/mg) and MG 0291 (7.28 μL/mg). The highest lignin contents were demonstrated in genotypes at 720 hai, which ranged between 10.79 and 12.17 μg/mg. Otherwise, the lowest levels of this compound were observed in MG 1207 (6.51 μg/mg) at 240 hai, and in MG 0291 (5.82 μg/mg) and Catuaí Vermelho IAC 144 (6.29 μg/mg) at 24 hai.

By analysing the soluble lignin contents at different times of collection, it was evidenced that MG 0291 genotype showed an increased content of this compound over the timeframe (Fig 4), unlike the MG 1207 and Catuaí Vermelho IAC 144. The lignin content in such genotypes at 24 and 240 hai were lower than observed at 120 and 480 hai, respectively. Between 480 and 720 hai, the same profile was verified in two genotypes: the last time of collection displayed a higher lignin content as compared to the previous ones.

## Discussion

Due to the environmental condition that benefits brown eye spot development in most coffee growing regions, besides the lack of resistant cultivars, this disease still representing a big challenge to Brazilian coffee growers, even after 100 years of introduction of desease in the country [16], or even 40 years after the first epidemic reports [17]. In this context, several efforts have been made by researchers worldwide to develop plants resistant to different *Cercospora* species; however, studies concerning *C. coffeicola* are still incipient. According to Nelson [18], environmental factors such as high altitudes, cloudiness, and high humidity benefit leaf brown eye spots development, which cause a high level of damage. Additionally, increases in the productive potential in coffee areas can lead to nutritional imbalance, resulting in a higher susceptibility of leaves to pathogen attack, which makes it difficult for breeders to select such plants in the field [3].

To face this challenge and seek to minimize environmental interference, our research group has been investigating the brown eye spot resistance under controlled conditions. Additionally, a collection of *C. coffeicola* strains sampled of *C. arabica* from all producing regions in Brazil were established to better understand plant-pathogen-environment relationships. Previously, it was reported the probability of fixing resistance genes to *C. coffeicola* in four accessions from the germplasm collection of Minas Gerais (GC-MG), as well as in the commercial cultivar Sarchimor MG 8840 [7]. In the present study, a mix of *C. coffeicola* strains was inoculated into 18 Brazilian commercial cultivars, an Arabica coffee clone, and 41 CG-MG accessions, to quantify the genetic resistance within the population and select genotypes to sustain the breeding program. After predicting the genotypic values of the studied materials, the most efficient way to select the severity data for brown eye spot was investigated.

Given the rejection of the null hypothesis for progeny variance (Table S1), this study aimed to point out strategies for early selection of genotypes that allow genetic gains in a short timeframe, seeking to further provide materials to the genetic breeding program for resistance in a field condition.

The accession MG 1207 Sumatra and cultivar Oeiras MG 6851 were considered partially resistant. On the other hand, accessions MG 0276, MG 0158 and cultivars Catiguá MG 2, Paraíso MG H 419-1 and IPR 103 were considered moderately susceptible. All the other genotypes studied were susceptible because they presented injured leaf area above 12.0%. In the present study, no genotype was classified as resistant. The existence of partially resistant phenotypes in our study indicated that severity to *C. coffeicola* is a quantitatively inherited character and presents an action conditioned by many genes with small individual effects and, consequently, it is highly influenced by environmental factors. In such cases, it is essential to use statistical methods based on estimates of genetic parameters, since heritability estimates enable both the development of more efficient selection strategies and predict the selection gain [19].

By using different approaches to analyze the outputs from severity data, it was demonstrated that only two genotypes (MG 1207 and Paraiso MG H 419-1) matched between both selection methods by considering five genotypes selected by AUCDP (MG 1207, MG 0321 and MG 0132, Paraiso MG H 419-1 and Asabranca) and five genotypes selected by the highest genotypic values and estimated heritability (MG 1207, Oeiras MG 6851, IPR 103, Paraíso MG H 419-1 and Catiguá MG 2). By estimating the heritability of each genotype, environmental effects are expected to be attenuated. Thus, it is noted that the genotypes MG 0321 and MG 0132 and Asabranca, selected by the AUCDP, presented a heritability below 70%, which indicates that the expression of the severity of brown eye spot in such materials presents a low genetic influence, and therefore responding majorly to environmental conditions [20]. Hence, it is not expected that such genotypes maintain their phenotypic behavior of low severity to brown eye spot over time, and this makes it difficult the experimental replication, which may be a false positive source of resistance. In this sense, only 20 genotypes showed high magnitude heritability (Table S1) within 60 evaluated ones. By considering the fixed model (Fig 2), only nine of these genotypes would have a satisfactory heritable fraction of genotypic variance - about 70% or more - (MG 1207, Paraiso MG H419-1, Asabranca, MG 0321, MG 0132, MG 0176, Sacramento MG1, MG 0280, MG 0380), if 20 genotypes of low severity to brown eye spot are selected. These results demonstrate that selection without observing the heritability of each accession individually may not be ideal, and this may lead to an inefficient method for breeding programs concerning resistance to *C. coffeicola*.

The genetic gain is inversely proportional to the selection intensity, which in the present study was 8.3% (five genotypes), thereby allowing higher efficiency in the next steps of selection [21]. It was observed that the accessions MG 1207, MG 0321 and MG 0132, and the commercial cultivars Paraiso MG H 419-1 and Asabranca presented a lower magnitude of their genotypic value to reduce the severity of brown eye spot, suggesting suitable possibilities of genetic progress in the sequence of evaluations (Table 1). Based on the severity of brown eye spot, the new predicted mean by the selection of these five genotypes was about 4.33, with a remarkable selection gain of 70% to reduce the severity (Fig 1). Accessions MG 1207 and MG 0132 refer to the Sumatra group, parental lines of the cultivar Mundo Novo; while the accession MG 0321 is the Hibrido de Timor UFV 432-09. The cultivar Paraiso MG H 419-1 represents the interbreeding between Catuaí Amarelo IAC 30 and the Híbrido de Timor UFV 445-46 [22]. Finally, the cultivar Asa Branca results from the interbreeding between Sarchimor 1668 and the World New 379-19 [23]. Therefore, these last three genotypes are resistant to the orange rust fungus of the coffee tree *H. vastatrix*.

Accuracy is a measure closely related to the precision of selection and holds the property of establishing the evaluation reliability and predict genotype genetic value based on its magnitude [20]. In the present study, it was observed that the assumption of four assessments resulted in values above or close to 99% (Table 1) by considering the estimated individual repeatability, which indicates a very high precision. It suggests that there are low absolute deviations between the true genotypic values and those estimated or predicted. Such outputs make it easier the achievement of expressive genetic gains. The efficiency of using four measurements as compared by using only one is about 1.23 or 23% for this character. These results corroborate with the number of measurements used in Arabica coffee trees to accomplish agronomic data [24]. By doubling this number, such as eight measurements, efficiency increases only 6%.

In this study, no significant differences were observed among the inoculated and non-inoculated genotypes of coffee plants with *C. coffeicola* regarding total soluble phenolic compounds and soluble lignin levels. Similar results have been reported elsewhere on this pathosystem [25, 26]. Since phenolic compounds and lignin are produced as a physical and chemical defense mechanism against pathogen attacks [27, 28, 29], it was expected that the inoculated genotypes would present higher levels of these metabolites as compared to non-inoculated ones.

The evidence that the levels of metabolites are not associated with coffee tree resistance to brown eye spot may be associated with phenylpropanoid pathway behaviour, from which phenolic compounds are synthesized via lignin monomers [30, 31]. By facing the stress, the genes involved in this pathway may have their expression altered to produce soluble phenolic compounds, which also present functions of defense and antioxidant properties, thus not necessarily being involved with lignin biosynthesis [32].

The crosstalk among the biosynthesis of phenolic compounds/lignin, growth, reproduction, and plant defense are other factors that can justify the fact that the inoculated and non-inoculated genotypes of coffee plants did not differ in metabolite levels. Plant defense and growth are negatively correlated, such as activation of defense processes that negatively affect plant growth and reproduction [29]. Thus, in this study, plants may have triggered other physiological processes rather than the accumulation of phenolic compounds and lignin (plant defense). As lignification is a process tightly controlled by various regulatory levels, it will only occur at the appropriate time and location of lignin deposition [32,31].

Leaf tissue necrosis results from the oxidation and polymerization of o-diphenols [33, 34]. In the present study, decreases in the content of phenolic compounds observed between 240 and 480 hai may be related to the use of such compounds for the synthesis of other products such as tannins and lignans, since the time of 480 hai is close to the period of symptoms manifestation of brown eye spot. On the other hand, increases in the content of phenolic compounds, verified between 480 and 720 hai, are characterized as a generic response in response to insects, fungi, or bacteria attacks [35]. Decreases in phenol content followed by successive increases were demonstrated elsewhere with the resistance of olive cultivars to *Verticillium dahliae* [36].

Lignin, on the other hand, plays an important role in plant growth and development, as it promotes increased cell wall rigidity and its metabolism is involved in the response to different biotic and abiotic stresses [37]. In this sense, environmental stresses change the content and composition of this compound in plants, which explains the difference observed in lignin content at the different times of collection in this study.

Based on the outputs presented in this study, the accession MG 1207 Sumatra was classified as partially resistant, which is in close agreement with the result previously demonstrated by Botelho et al [7]. This genotype can substantially contribute to the development of a new cultivar to reduce the use of pesticides. Herein, it is also evidenced that the assumption of four assessments for the severity of brown eye spot is enough for parameters accuracy and efficiency, leading to expressive genetic gains. Finally, it is suggested that the levels of lignin and phenolic compounds are not associated with the resistance of coffee genotypes to brown eye spot.

## Acknowledgments

The authors wish to thank National Counsel of Technological and Scientific Development-CNPq, the Brazilian Consortium Coffee Research and Development, the National Institute of Coffee Science and Technology (INCT Café/CNPq), and the Foundation for Research Support of the State of Minas Gerais (FAPEMIG) for their financial support. The funders had no role in study design, data collection and analysis, decision to publish, or preparation of the manuscript.

## Authors’ contributions

All authors contributed to the study conception and design. Material preparation, data collection and analysis were performed by Juliana B. Ramos; Mario L.V. de Resende; Deila M. dos S. Botelho; Renata C. M. Pereira; Tharyn Reichel; André A. F. Balieiro; Gustavo P. Botega^2^; Juliana C. de R. Abrahão. The first draft of the manuscript was written by Juliana B. Ramos and all authors commented on previous versions of the manuscript. All authors read and approved the final manuscript.

---

**S1 Table:**
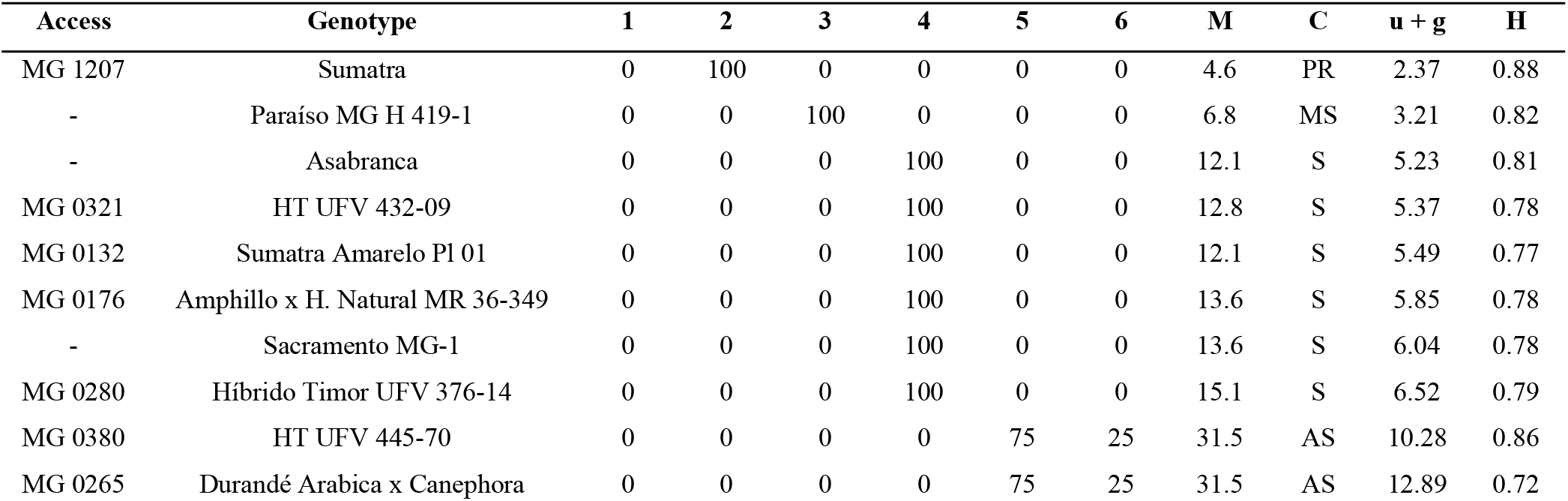
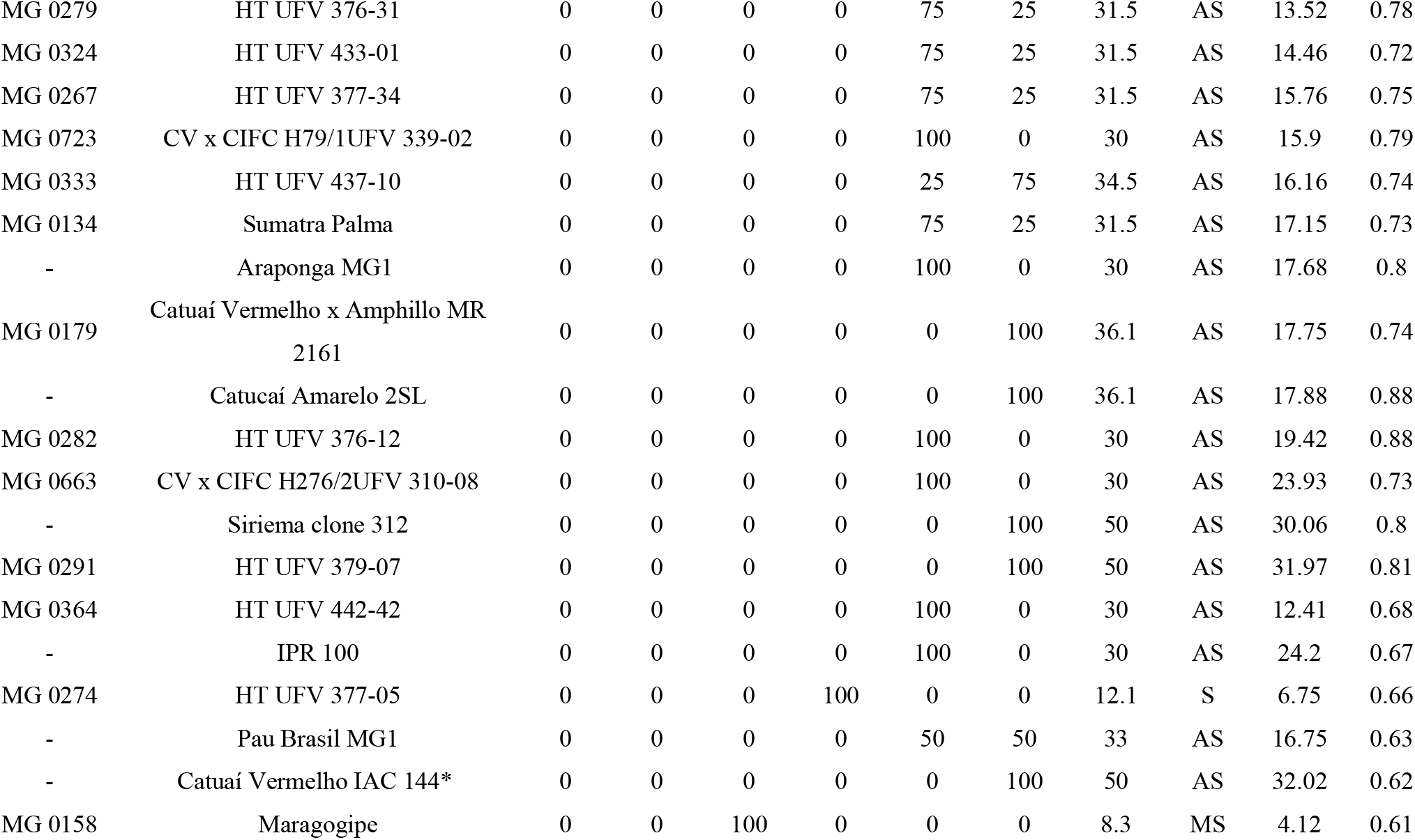
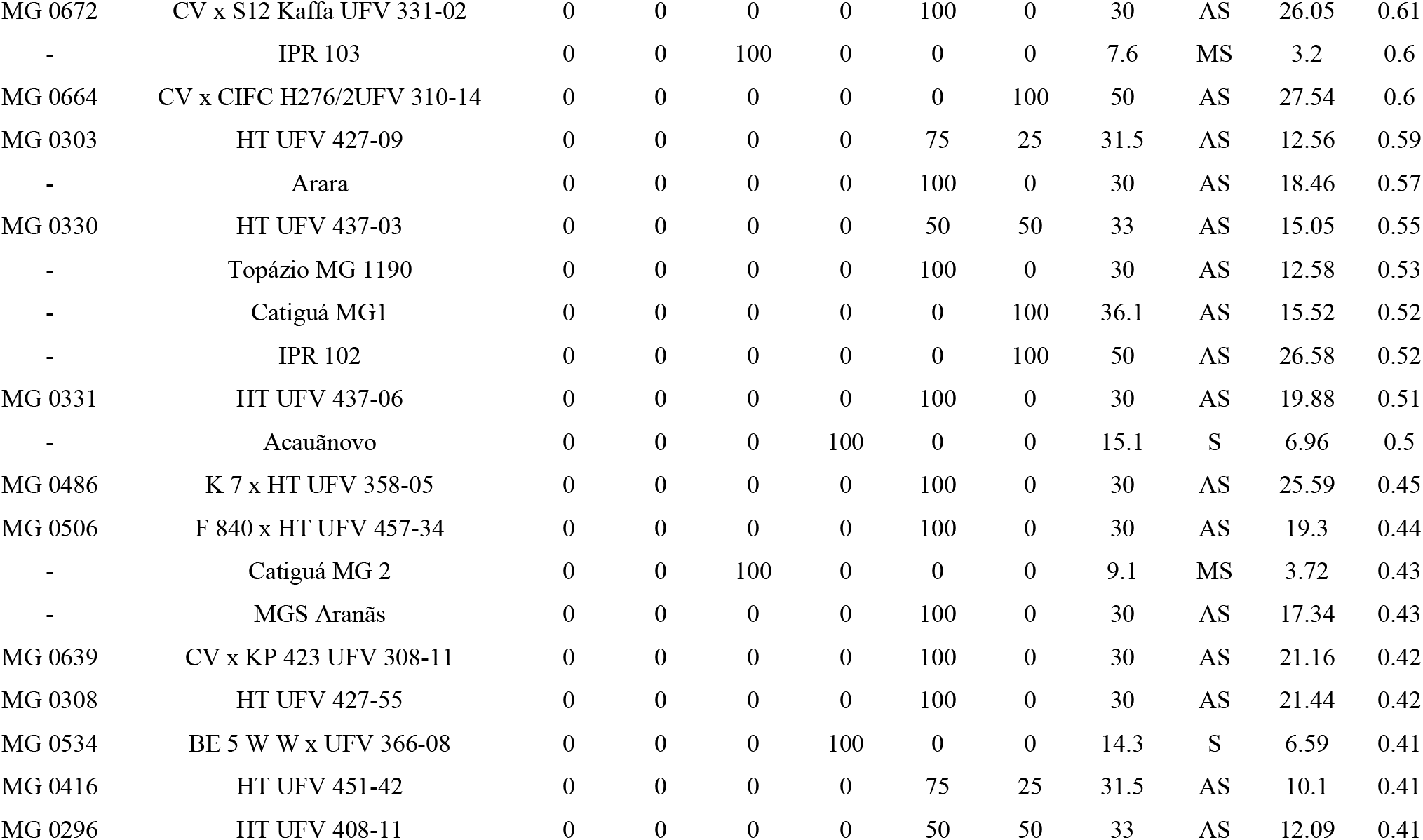
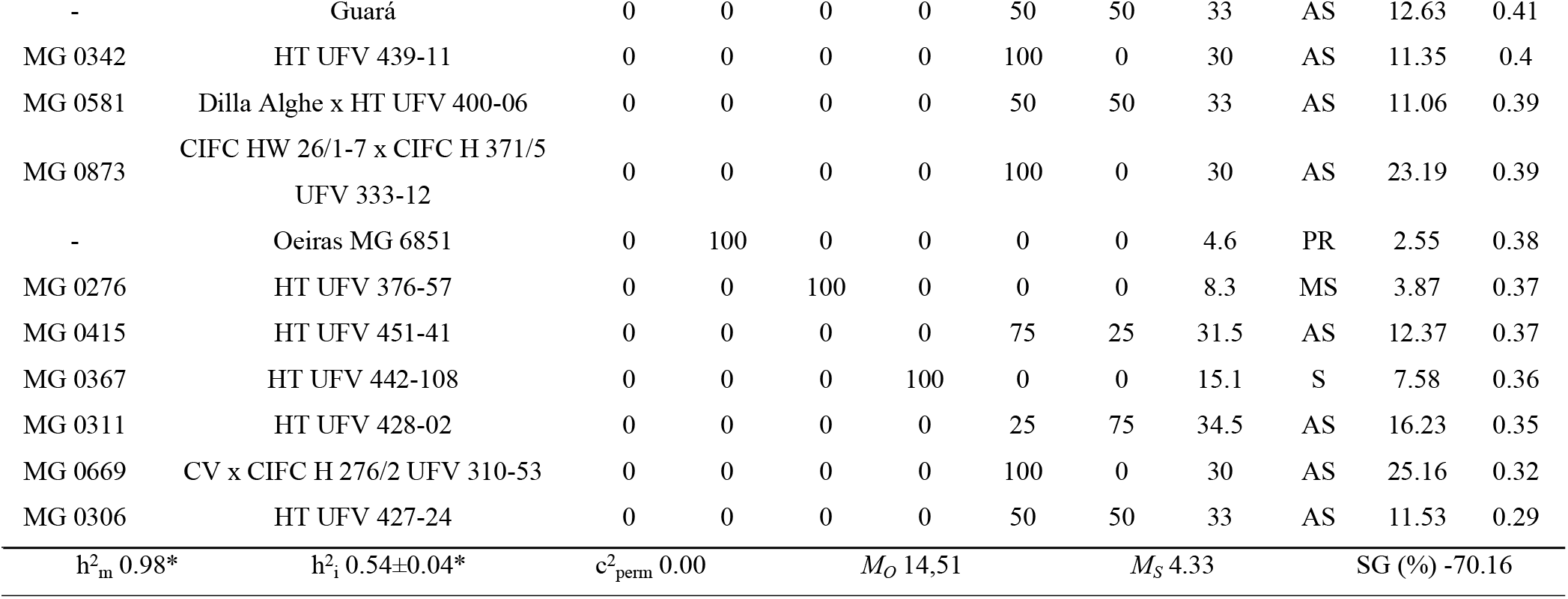
Genotypes and their respective distribution of leaf lesions frequency into six classes, class 1: 0.1-3.0; class 2: 3.1-6.0; class 3: 6.1-12.0; class 4: 12.1-18.0; class 5: 18.1-30.0; class 6: 30.1-50% of the leaf surface affected by brown eye spot, phenotypic mean of the genotypes (M), classification (C) of the accessions based on the phenotypic mean, predicted additive breeding values (u + g), heritability (H) of genotypes, genetic parameters, general mean and estimated gain relative to the severity of brown eye spot in 5 selected genotypes. PR: partially resistant, MS: moderately susceptible, S: susceptible, AS: highly susceptible. *Cultivar used as a control. CV – Catura Vermelho; HT – Híbrido de Timor h^2^_m_: heritability of genotype averages. h^2^_i_: broad-sense heritability individually for each plot, or the total genotypic effects. c^2^_perm_: coefficient of determination of the permanent environmental effects. M_O_: Genotypic mean of 60 genotypes before selection. M_*S*_ Mean of the five selected genotypes. SG: gain selection.

